# Reconciling a Model of Core Metabolism with Growth Yield Predicts Biochemical Mechanisms and Efficiency for a Versatile Chemoautotroph

**DOI:** 10.1101/717884

**Authors:** Jesse McNichol, Stefan M. Sievert

## Abstract

Obligately chemoautotrophic *Campylobacteria* dominate productivity in dark, sulfidic, and oxygen-depleted environments. However, biochemical mechanisms underlying their growth remain poorly known, limiting understanding of their physiology, ecology, and biogeochemical impact. In this study, we used comparative genomics, conceptual modeling of core metabolism, and chemostat growth yields to derive a model of energy conservation consistent with experimental data for the versatile chemoautotroph *Sulfurimonas denitrificans*. Our model rests on three core mechanisms: Firstly, to allow electrogenic sulfur-based denitrification, we predict that the campylobacterial-type sulfur oxidation enzyme complex must donate electrons to the membrane quinone pool, possibly via a sulfide:quinone oxidoreductase. Secondly, to account for the unexpectedly low growth efficiency of aerobic sulfur oxidation compared to denitrification, we posit the high-affinity campylobacterial-type cbb_3_ cytochrome c oxidase has a relatively low H+/e− of 1, likely due to a lack of proton pumping under physiological conditions. Thirdly, we hypothesize that reductant for carbon fixation by the reverse tricarboxylic acid cycle is produced by a non-canonical complex I that reduces both ferredoxin and NAD(P)H. This complex is conserved among related *Campylobacteria* and may have allowed for the radiation of organisms like *S. denitrificans* into sulfur-rich environments that became available after the great oxidation event. Our theoretical model has two major implications. Firstly, it sets the stage for future experimental work by providing testable hypotheses about the physiology, biochemistry, and evolution of chemoautotrophic *Campylobacteria*. Secondly, it provides constraints on the carbon fixation potential of chemoautotrophic *Campylobacteria* in sulfidic environments worldwide by predicting theoretical ranges of chemosynthetic growth efficiency.

**Significance:** Chemoautotrophic *Campylobacteria* are abundant in many low-oxygen, high-sulfide environments where they contribute significantly to dark carbon fixation. Although the overall redox reactions they catalyze are known, the specific biochemical mechanisms that support their growth are mostly unknown. Our study combines conceptual modeling of core metabolic pathways, comparative genomics, and measurements of physiological growth yield in a chemostat to infer the most likely mechanisms of chemoautotrophic energy conservation in the model organism *Sulfurimonas denitrificans*. The hypotheses proposed herein are novel, experimentally falsifiable, and will guide future biochemical, physiological, and environmental modelling studies. Ultimately, investigating the core mechanisms of energy conservation will help us better understand the evolution and physiological diversification of chemoautotrophic *Campylobacteria* and their role in modern ecosystems.

## Introduction

At deep-sea hydrothermal vents, chemoautotrophic microbes support entire food webs by fixing inorganic carbon into biomass (1, 2). Chemoautotrophs from the class *Campylobacteria* often dominate the microbial communities at deep-sea vents and in other sulfidic environments, coupling the oxidation of hydrogen and reduced sulfur compounds to the reduction of nitrate, oxygen and sulfur to support autotrophic CO_2_ fixation via the reverse tricarboxylic acid (rTCA) cycle (3–6). Aside from their important environmental role, the *Campylobacteria* are of interest from an evolutionary perspective. Long known as the epsilon subdivision of the *Proteobacteria*, this group has been recently reclassified as the class *Campylobacteria* in the new phylum *Campylobacteraeota* to reflect their divergent evolutionary heritage (6). Recent data suggest that *Campylobacteria* possess a number of unique biochemical characteristics that may be essential for their success as chemoautotrophs in sulfidic environments (7–11). Despite these data, we still lack information on the specific mechanisms of chemoautotrophic energy conservation for *Campylobacteria*. One major reason for this knowledge gap is that their fastidious microaerophilic/anaerobic growth phenotype makes them difficult to manipulate experimentally.

In this study, we addressed these unknowns by developing an experimentally-consistent theoretical model of core metabolism for the campylobacterial chemoautotroph *Sulfurimonas denitrificans* (12, 13). This organism is metabolically-flexible, capable of using sulfide, sulfur, thiosulfate, and hydrogen as electron donors as well as oxygen or nitrate as terminal electron acceptors. In addition, *S. denitrificans* is comparatively well-studied from a physiological and genomic perspective among chemoautotrophic *Campylobacteria* (12, 14–19). This makes *S. denitrificans* an ideal candidate to investigate multiple different energy-conserving processes and their potential biochemical underpinnings.

To construct our conceptual model, we first identified potential pathways that support energy conservation in *S. denitrificans* using comparative genomics and biochemical knowledge from related organisms (15, 20). This resulted in a number of possible mechanisms and/or proton translocation stoichiometries for core energy-conserving protein complexes. Next, biomass yields measured in continuous culture (this study, (14) were compared with yields predicted from these putative models to determine the most parsimonious biochemical mechanism for chemoautotrophic growth in *S. denitrificans*. Below, we discuss the logic of model construction and how it led us to predict novel mechanisms of energy conservation. We also propose direct experimental tests and discuss the implications of these hypothetical mechanisms for understanding the bioenergetics, evolution, and niche differentiation of diverse chemoautotrophic *Campylobacteria*.

## Results and Discussion

We developed a theoretical model of energy conservation coupled to carbon fixation by the rTCA cycle for the chemoautotrophic bacterium *Sulfurimonas denitrificans* (Table 1; Figure 1). To determine the most parsimonious model, we compared predicted relative growth yield ratios (RGY) of hypothetical mechanisms to substrate growth yield ratios measured in a chemostat (Table 2; 14). Using this approach, we eliminated implausible mechanisms and converged on a model that accurately reproduced RGY in *S. denitrificans* under three distinct growth conditions (Tables 1 and 2; see discussion below and in SI). Our model also allowed us to constrain the chemosynthetic growth efficiency (CGE) of *S. denitrificans*. CGE is a measure of the overall efficiency of metabolism, representing the fraction of total electrons from electron donors used to fix CO_2_ (equivalent to 1 - y in reference (21); Table 1). While the biochemical reactions that we inferred have explicit stoichiometry, they could not directly be used to calculate CGE. This is because CGE is also dependent on the amount of proton-motive force (PMF) required for growth in addition to carbon fixation (e.g. ATP/PMF for other assimilatory/biosynthetic reactions or for motility/transport). This quantity, hereafter referred to as “non-rTCA-PMF” was also inferred from our model and RGY data as follows.

**Table 1.** Hypothetical metabolic reactions for S.denitrificans inferred from comparative genomics and pathway logic, and calculated maximum chemosynthetic growth efficiency. See text for derivation and references.

| <u>Half reactions</u> |  |  |  |
| --- | --- | --- | --- |
| Dissimilatory oxidation |  | Description | Equation stoichiometry |
| 1 (S2O3-OX) | | Thiosulfate oxidation by SOX linked to menaquinone pool | $S2O3^{2-} + 4 MK + 5 H2O + 8 H^+ (p) \Rightarrow 2 SO4^{2-} + 10 H^+ (p) + 4 MKH2$ |
| 2 (H2-OX) | | Hydrogen oxidation linked to menaquinone pool | $H2 + 2 H^+ (c) + 1 MK \Rightarrow 2 H^+ + 1 MKH2$ |
| Dissimilatory reduction |  | Description | Equation stoichiometry |
| 3 (QcrABC) | | Electron transport to cytc by QcrABC (overall reaction) | $MKH2 + 2 H^+ (c) \Rightarrow MK + cytc2^+ + 2 H^+ (p)$ |
| 4 (NO3-RED) | | Nitrate reduction by Nap complex (with QcrABC) | $NO3^- + MKH2 + 2 H^+ (c) + 2 H^+ \Rightarrow 2 H^+ (p) + H2O + NO2^- + MK$ |
| 5 (NO2-RED) | | Nitrite reduction by Nir complex | $2 H^+ + NO2^- + cytc2^+ \Rightarrow H2O + cytc3^+ + NO$ |
| 6 (NO-RED) | | Nitric oxide reduction by NorBC complex | $2 H^+ + 2 NO + 2 cytc2^+ \Rightarrow 2 cytc3^+ + 1 N2O + 1 H2O$ |
| 7 (N2O-RED) | | Nitrous oxide reduction by Nos complex (with QcrABC) | $N2O + MKH2 + 2 H^+ (c) + 2 H^+ \Rightarrow N2 + MK + 2 H^+ (p) + H2O$ |
| 8 (O2-RED) | | Oxygen reduction by cbb3 (PMF produced by chemical protons) | $O2 (p) + 4 H^+ (c) + 4 cytc2^+ \Rightarrow 2 H2O + 4 cytc3^+$ |
| <u>Reverse electron transport</u> |  | Description | Equation stoichiometry |
| 8 (CPLX1) | | NADPH and ferredoxin reduction via EPSI-type Complex I | $NAD(P)^+ + 2 Fd(ox) + 8 H^+ (p) + 2 MKH2 \Rightarrow NAD(P)H + 11 H^+ + 2 MKH2 + 2 Fd(red)$ |
| 9 (CFIX-rTCA) | | Overall reaction for rTCA cycle | $4 CO2 + 4 Fd(red) + 2 NAD(P)H + MKH2 + 3 ATP \Rightarrow 1 Oxaloacetate + 4 Fd(ox) + 2 NAD(P)^+ + MK + 3 ADP$ |
| <u>Coupled reactions and predicted chemosynthetic growth efficiency (CGE)</u> |  |  |  |
| Pathway | Description | Reactions | Theoretical CGE* |
| T/N | S2O3-dependent denitrification | 1 & 3-7 | 0.11 - 0.14 |
| T/O | S2O3-dependent oxic respiration | 1 & 8 | 0.20 - 0.25 |
| H/N | H2-dependent denitrification | 2 & 3-6 | 0.23 - 0.28 |
| H/O | H2-dependent oxic respiration | 2 & 8 | 0.31 - 0.38 |

\*CGE was calculated by estimating the amount of proton-motive force/ATP required for biosynthesis in addition to that required for carbon fixation. This was in turn estimated from experimentally measured yield ratios between H/N and T/N conditions (Table 2; see text and SI for further explanation).

**Figure 1:**
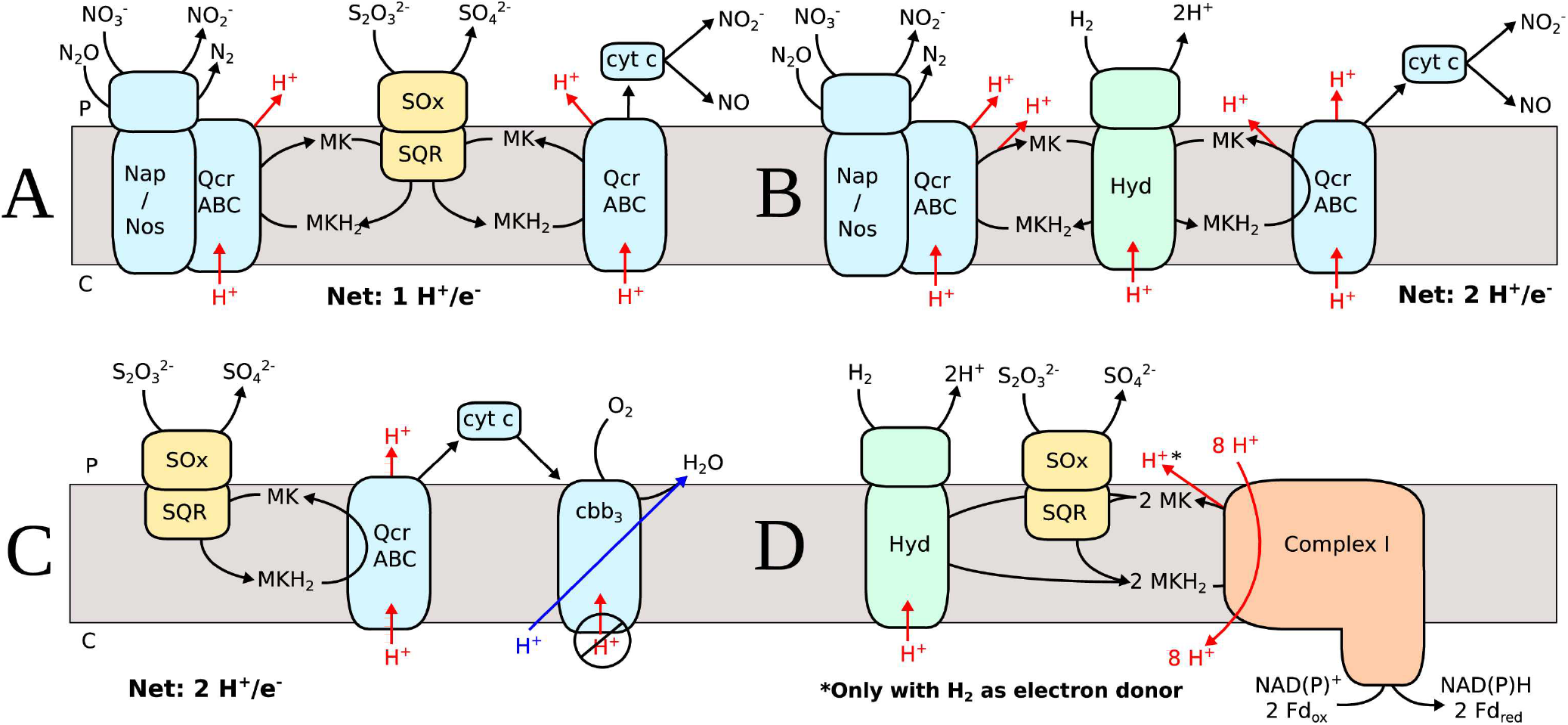
Hypothetical model of electron transport and energy conservation in *Sulfurimonas denitrificans* during: A) Thiosulfate-dependent denitrification, B) Hydrogen-dependent denitrification, C) Thiosulfate-dependent aerobic respiration and, D) Reversed electron transport to produce reductant for the rTCA cycle. See text for explanation. Black arrows indicate electron transport, red arrows contributions to the proton-motive force and blue arrows are “chemical protons” used during reduction of O_2_ to H_2_O. Nap=Nap nitrate reductase complex; Sox=Sox sulfur oxidation complex; SQR=Sulfide:quinone oxidoreductase; Hyd=Hydrogenase; QcrABC=Complex III; MK/MKH_2_=Oxidized/reduced menquinone; cyt c=Soluble cytochrome c; cbb_3_=high-affinity cytochrome c oxidase; Fd=soluble ferredoxin; P=Periplasm; C=Cytoplasm.

**Table 2.** Predicted relative growth yields compared with growth yields measured from chemostat experiments

| Growth Condition | Modelled RGY<sup>†</sup> | Y<sub>SO4</sub><sup>*</sup> (this study) |  | Y<sub>S2O3 / 2</sub><sup>*</sup> (Timmer Ten-Hoor, 1981) |  |  |
| --- | --- | --- | --- | --- | --- | --- |
|  |  | AFDW (n=3) | CC (n=3) | DW | CC | CP |
| S<sub>2</sub>O<sub>3</sub><sup>2-</sup>/NO<sub>3</sub><sup>-</sup> |  | 0.89 ± 0.14 | 0.67 ± 0.14 | 2.58 | 1.18 | 0.89 |
| S<sub>2</sub>O<sub>3</sub><sup>2-</sup>/O<sub>2</sub> |  | 1.49 ± 0.20 | 0.91 ± 0.12 | 3.87 | 2.00 | 1.95 |
| Relative growth yield (RGY) | 1.55 - 2 ‡ | 1.67 | 1.36 | 1.50 | 1.70 | 2.20 |

| Growth Condition | Modelled RGY<sup>†</sup> | Y<sub>NO3</sub><sup>*</sup> (this study) |  | * = Biomass yield per mole substrate<br>AFDW = Ash-free dry weight (g • M<sup>-1</sup>)<br>CC = Cell carbon (g C • M<sup>-1</sup>)<br>DW = Dry weight (g • M<sup>-1</sup>)<br>CP = Cell protein (g protein • M<sup>-1</sup>) |
| --- | --- | --- | --- | --- |
|  |  | AFDW (n=3) | CC (n=3) |  |
| S<sub>2</sub>O<sub>3</sub><sup>2-</sup>/NO<sub>3</sub><sup>-</sup> |  | 1.01 ± 0.22 | 0.76 ± 0.21 |  |
| H<sub>2</sub>/NO<sub>3</sub><sup>-</sup> |  | 2.45 ± 0.73 | 1.75 ± 0.31 |  |
| Relative growth yield (RGY) | 2 - 3.33 ‡ | 2.43 | 2.30 |  |

† = Predicted growth yield based on fixation of one oxaloacetate molecule (Reaction 9 in Table 1), and ranges (‡) are as a function of the fraction of “non-rTCA-ATP” used by the cell (see text, supplemental text, and Fig S1 and S2 for further explanation).

Assuming that the amount of non-rTCA-PMF is identical between growth conditions, both RGY ratios (H/N : T/N and T/O : T/N) will approach 2 as the amount of non-rTCA-PMF relative to that needed for carbon fixation increases (Figures S1 and S2). Using this information, we inferred the proportion of non-rTCA-PMF and a range of theoretical CGE (Table 1; see SI for more detail). These theoretical values were very close to chemostat yield measurements previously reported for *S. denitrificans* and other members of the genus *Sulfurimonas* (0.167 and 0.098 - 0.148, for microaerobic and denitrifying growth respectively; summarized in 21). Although CGE will ultimately be dependent on the fraction of non-rTCA-PMF required under a given growth condition, our theoretical approach allows us to calculate explicit bounds on CGE and therefore constrain maximal primary productivity in ecosystems dominated by chemoautotrophic *Campylobacteria*. Below, we describe the three biochemical mechanisms predicted from our approach and their implications for understanding the evolution and ecology of chemoautotrophic *Campylobacteria*.

### Thiosulfate-Dependent Denitrification

*S. denitrificans* was isolated in a nitrate-limited chemostat with thiosulfate as the sole electron donor (12). We initially based our model of sulfur oxidation on previously-validated biochemical mechanisms for the SOx sulfur oxidation system, whereby it donates electrons at the level of soluble cytochrome c (22, 23). However, this canonical SOx mechanism cannot support electrogenic thiosulfate-dependent denitrification since electrons would have to be transferred endergonically (“uphill”) to the quinone-dependent Nap/QcrABC nitrate reductase supercomplex (24) and the nitrous-oxide reducing Nos/QcrABC supercomplex (25). Although the intervening enzymatic steps in denitrification (nitrite and nitric oxide reduction) likely accept electrons at the level of cytochrome c, these complexes are thought to be non-electrogenic, meaning no energy could be generated with thiosulfate/nitrate in the absence of QcrABC (24–26). We therefore concluded that the most likely explanation is the existence of an alternative “campylobacterial-SOx” system that donates electrons from sulfur oxidation to the membrane quinone pool (Fig 1a).

Biochemical data tentatively support the existence of a membrane-bound SOx complex in chemoautotrophic *Campylobacteria* and other sulfur-oxidizing bacteria. In *Sulfurovum* NBC37-1, both membrane-bound and soluble protein fractions are necessary for thiosulfate oxidation (8); similar data have been reported for the alphaproteobacterium *Starkeya novella* (27, 28). What protein(s) potentially serve as a membrane anchor for the SOx system in *S. denitrificans* remains unknown. Yamamoto et al. (2010) identified several proteins involved in thiosulfate oxidation that have putative membrane-binding motifs, including SoxH, a cytochrome c, and a sulfide:quinone oxidoreductase (SQR)-like protein. SQR proteins are an attractive candidate for an campylobacterial-SOx anchor as they are known to be membrane-bound and capable of sulfide-dependent quinone reduction in *S. denitrificans* and other bacteria (17, 29). Consistent with its possible role in thiosulfate oxidation, RT-qPCR shows that a putative SQR in *S. denitrificans* (Suden_0619) is highly expressed in the absence of sulfide (Figs S3 and S4).

By including the campylobacterial-SOx complex, our model was able to closely reproduce experimental RGY data between hydrogen and thiosulfate-based denitrification conditions (Table 2) (16). We predict higher growth yields with hydrogen because the membrane-spanning hydrogenase in *S. denitrificans* (15) abstracts protons from the cytoplasm during hydrogen-dependent reduction of menaquinone. When menaquinol is reoxidized, protons are released into the periplasm (Fig 1b), thus generating a proton-motive force via a redox-loop mechanism (20). In contrast, periplasmic sulfur-oxidizing enzymes likely do not span the membrane (8, 30). Therefore, thiosulfate-dependent quinone reduction itself is probably electroneutral, although a proton-motive force is still generated by the action of complex III (QcrABC). QcrABC oxidizes quinol and either transfers electrons to soluble cytochrome c or acts together as a supercomplex with the Nap/Nos complexes that reduce nitrate and nitrous oxide respectively (Fig 1). The overall effect is that QcrABC makes electron transport to nitrite, nitric oxide and nitrous oxide an electrogenic process (25). Complex III will also conserve energy with hydrogen oxidation, which is in addition to the energy from the aforementioned redox loop.

### Aerobic Respiration vs. Denitrification

*S. denitrificans* is able to grow microaerobically with thiosulfate, which yields ~ 60 % more biomass than with thiosulfate/nitrate (Table 2; (14)). Microaerobic respiration is most likely sustained by a high-affinity cytochrome c oxidase (cbb_3_; complex IV) as no other putative oxygen-reducing complexes are present in the genome. The number of protons pumped by cbb_3_ is controversial, but some have suggested that it has the potential to equal low-affinity forms of complex IV (i.e. mitochondrial aa_3_; (31)). However, modelling *S. denitrificans’* growth using the same proton-pumping stoichiometry as aa_3_ coupled to complex III (3 H^+^/e^−^; (20)) presented a problem - both CGE and the RGY between aerobic and denitrifying conditions were far higher than observed in this study (Tables 1 and 2) and in (14).

To reconcile our model with growth yield data, the most parsimonious explanation is that complex IV in *S. denitrificans* has an overall H^+^/e^−^ of 1 (in contrast to the canonical aa_3_ complex which both pumps protons and consumes cytoplasmic protons for a total of a H^+^/e^−^ of 2). This could occur if either no protons are pumped during aerobic respiration or if the protons used in the reduction of oxygen (hereafter “chemical protons”) are abstracted from the periplasm instead of the cytoplasm as is known for the aa_3_ complex (Fig 1c). We believe the former explanation is more likely, as experiments have shown that cbb_3_-type cytochrome c oxidases behave differently during *in vitro* biochemical assays (32) and cannot sustain proton pumping at physiologically-relevant membrane potentials ((33); see also SI). This is due to the fact that high-affinity cytochrome c oxidases use more free energy in binding their substrate, resulting in less energy available to pump protons against a membrane gradient (33). We also note several differences exist between the “campylobacterial-type cbb_3_” found in *S. denitrificans* and other bacteria. For example, the *fixGHIS* genes are missing (15), as well as a number of histidine residues conserved elsewhere (34) - one of which has been shown to have an important catalytic role (35).

### Reversed Electron Transport

Another pathway essential to the growth of *S. denitrificans* is the chemoautotrophic fixation of CO_2_ via the rTCA cycle (36). In order to support carbon fixation, reduced ferredoxin is required as an electron donor for reductive carboxylation of succinyl-CoA to alpha-ketoglutarate and acetyl-CoA to pyruvate (36, 37). However, how reduced ferredoxin is produced in chemoautotrophic *Campylobacteria* is unclear (38). Sievert et al. (2008) suggested that hydrogen-dependent electron bifurcation could generate both NAD(P)H and reduced ferredoxin. However, since this mechanism depends on hydrogen, it cannot reduce ferredoxin with sulfur compounds as an electron donor. In addition, the soluble hydrogenase hypothesized to be responsible by Sievert et al. (2008) is not expressed at high levels in mRNA under a variety of conditions (Figs S3 and S4; (19)). No known electron-bifurcating complexes are present in the *S. denitrificans* genome (38), nor is there evidence for complexes that couple endergonic ferredoxin reduction to the proton motive force (i.e. the Rnf complex; (39)).

A more plausible mechanism for ferredoxin production in *Campylobacteria* was uncovered by Weerakoon and Olson (2008) in a study of the heterotroph *Campylobacter jejuni*. These authors report that the NADH:ubiquinone oxidoreductase (complex I) in *C. jejuni* does not accept electrons from NADH. Instead, reduced flavodoxin serves as the physiological electron donor during heterotrophic growth. Since flavodoxin can substitute for ferredoxin (40), such a non-canonical complex I run in reverse has the potential to reduce ferredoxin. Weerakoon and Olson (2008) demonstrated that this unusual mechanism is due to differences in the NuoE/F subunits that interact with NADH in canonical complex I. They also note that *S. denitrificans* and other chemoautotrophic *Campylobacteria* possess non-canonical NuoEF subunits and two copies of NuoG. Based on currently available genomes, this non-canonical complex I subunit composition is universally conserved among the chemoautotrophic genera *Sulfurimonas* and *Sulfurovum* (15, 41–45)(NZ_CP011308). Considering this significant and evolutionarily-conserved structural difference in complex I among chemoautotrophic *Campylobacteria*, it appears likely to have an essential role in core metabolism. However, NAD(P)H is also needed as a reductant for the rTCA cycle, and the mechanism demonstrated by Werakoon and Olson does not involve NAD(P)H. In *Hydrogenobacter thermophilus,* a “plant-like” ferredoxin:NADP+ reductase transfers electrons from ferredoxin to NADPH (46), whereas in *Nitrospina gracilis* genomic evidence suggests two distinct complexes may be specialized for NAD(P)H or ferredoxin reduction, respectively (47). However, there is no evidence for either of these mechanisms in *S. denitrificans*.

How then does *S. denitrificans* then produce NAD(P)H? An important clue comes from comparison of the protein replacing NuoF in *S. denitrificans* (Suden_1824) to known electron-bifurcating complexes. We note that Suden_1824 is homologous to the NfnB subunit of the NfnAB complex of *Thermotoga maritima* (33% AA ID; 88% coverage; Fig S5) which transfers electrons from NADPH to both reduced ferredoxin/NADH during flavin-based electron bifurcation (48); this protein was also recently shown to bind NADPH and probably also ferredoxin (49). Relevant residues for binding the phosphate group of NADP(H) (Tyr310, Arg311-312 in *T. maritima*) and cysteines ligating the iron-sulfur clusters that putatively pass electrons to ferredoxin are conserved in Suden_1824, as well as most residues involved in binding FAD ((49); Fig S5). Therefore, we propose that the function of complex I in *S. denitrificans* is to reduce NAD(P)+ and ferredoxin simultaneously during reversed electron transport. In this hypothetical reaction, 4 electrons derived from two menaquinol molecules reduce one molecule of NAD(P)+ and two molecules of ferredoxin via an electron-bifurcation mechanism (Fig 1d). We hypothesize that this endergonic reaction is driven by the transport of eight protons from the periplasm into the cytoplasm. A similar mechanism has been proposed in anammox bacteria to produce both NADH and formate simultaneously based on a similar non-canonical complex I structure (50). We also note recent work showing that modifications to this core bioenergetic complex are commonly found in other taxa, presumably to accommodate different electron carriers or increase proton-pumping efficiency (51).

If our hypothesis is true, then a site for electron bifurcation must exist in *S. denitrificans’* complex I. One possible location is in the NuoG subunit, which contains an iron-sulfur cluster (N7) that is separated from the electron transfer pathway and thought to be an evolutionary relic in organisms such as *E. coli* (52). However, NuoG in *Campylobacteria* and *Aquifex aeolicus* contain four conserved cysteine residues that have been shown to ligate an additional iron-sulfur cluster in *A. aeolicus (34, 53, 54)*. This additional cluster may connect the N7 “relic” iron-sulfur cluster to electron transport, allowing electron bifurcation to occur at N4. In addition, *S. denitrificans* and close relatives contain two distinct copies of NuoG, both of which are expressed at a similar level in transcriptomes of *S. denitrificans* (19). These two copies have similar sequences (36 % amino acid identity), but differ in terms of length. The short subunit (Suden_1823) is “missing” two main regions (Fig S6). The first (AA 466-532 of Suden_1822) contains no obvious functional motifs. The second missing region (AA 175-204 of Suden_1822) is found between two cysteine residues that putatively bind an additional iron-sulfur cluster (34). Without this stretch of sequence, the distance between these two cysteines is shortened to 29 amino acids which is very similar to the *A. aeolicus* NuoG (53). Therefore, Suden_1823 may be a potential location for electron bifurcation. What then could be the function of the other NuoG subunit? Perhaps it is involved in shuttling electrons to the non-canonical NuoF subunit, or maybe it is itself involved in binding ferredoxin (55).

The ability to use complex I to produce cellular reductant would have several advantages for *S. denitrificans*. Since this mechanism is dependent on the proton motive force, it would be linked to (and regulated by) energy generation processes and independent of the electron donor (Fig 1d). Additionally, it provides both reductants in a 1:1 ratio. Since the rTCA cycle uses electrons from ferredoxin and NAD(P)H in a 1:1 ratio if fumarate reduction occurs with quinol as the electron donor (15, 56), this balanced stoichiometry would be important for balancing the redox state of the cell.

### Environmental Implications

Our model and experimental data show how different electron donor/acceptor combinations greatly affect the CGE of *S. denitrificans*. For example, under denitrifying conditions, the use of hydrogen allows for an approximate doubling of CGE versus the use of thiosulfate (Table 1). Similarly, since aerobic respiration pumps twice as many protons as denitrification, it is the most efficient electron-accepting process for *S. denitrificans*. Our model therefore predicts that hydrogen-dependent aerobic respiration is the most efficient dissimilatory pathway for *S. denitrificans* - approximately 3 times greater than growth under the T/N condition.

Future work should include both biochemical and physiological studies to test our hypotheses (Table S1). These physiological experiments may be challenging to conduct. For example, measuring the CGE of *S. denitrificans* with hydrogen and oxygen as a donor/acceptor pair is complicated by the obligate microaerobic nature of *S. denitrificans.* Providing a continuous, non-limiting supply of low levels of oxygen while simultaneously measuring low levels of consumption will require development of novel experimental procedures. With regards to biochemical validation, although we have directly identified some biochemical targets (e.g. complex I and IV), others are currently unknown (e.g. the posited membrane anchor of the SOX complex). To address this uncertainty, techniques such as transposon-mutagenesis sequencing (57) may allow for the identification of putative target genes without relying on genetic approaches that are difficult to apply to organisms such as *S. denitrificans*.

## Conclusions

Previous work has used thermodynamics to estimate the amount of energy available for microbial life at hydrothermal vents and thus their potential for generating new organic carbon (58–60). Our recent field measurements have complemented these theoretical approaches by directly measuring the rates and efficiency of carbon fixation by chemoautotrophs in hydrothermal fluids under natural temperature/pressure conditions (2, 61). Here, we aimed to better understand the mechanistic basis behind these processes by applying a theoretical framework relying on advances in genomics and recent discoveries of bioenergetic mechanisms. Equally important, we also used classical cultivation and growth yield techniques that were last applied to *S. denitrificans* nearly 45 years ago (12, 14). It was only by incorporating these physiological data that we could justify our predictions of novel mechanisms putatively underlying the growth of *S. denitrificans* and similar chemoautotrophic *Campylobacteria.* Not only are our hypotheses parsimonious and consistent with experimental data, they also provide a theoretical framework for designing biochemical experiments (Table S1) and modelling natural ecosystems.

In a broader sense, studying the function and mechanisms of core energy-conserving enzymes will also help understand the evolutionary trajectory of the *Campylobacteria*. This point can be best illustrated by the non-canonical complex I found among the *Campylobacteria*. In contrast to *S. denitrificans,* deep-branching representatives possess a much simpler complex I that completely lacks the NuoEFG subunits that probably interact with ferredoxin and/or NAD(P)H (7, 62). Such a truncated complex resembles so-called energy-converting hydrogenases (Ech) that are thought to be the evolutionary precursors of complex I (63). Since this form of complex I is only found in hydrogen-dependent and oxygen-sensitive taxa, it may represent the ancestral form of this core bioenergetic complex in *Campylobacteria*. In contrast, for oxygen-tolerant organisms such as *S. denitrificans* that do not rely on hydrogen for growth, complex I is larger and contains additional subunits that were probably acquired at a later time. We speculate that these additional subunits may have allowed for the adaptive radiation of chemoautotrophic *Campylobacteria* into oxic, sulfide-rich niches that became available during the rise of atmospheric oxygen on the ancient Earth (6, 64). Therefore, by studying the physiology and biochemistry of chemoautotrophic *Campylobacteria,* we can gain insight into key evolutionary changes that occurred in tandem with the biogeochemical evolution of the biosphere.

## Materials and Methods

### Development of conceptual model

Based on knowledge from the initial genome characterization of *S. denitrificans* (15), curation of core energy-conserving metabolic reactions was carried out to reproduce *S. denitrificans’* observed physiology (14–16). Reactions were based on Fig. 2 from Sievert et al. (2008) with the following modifications. Cytochrome cbb_3_ uses only oxygen (not nitric oxide as proposed in Sievert et al. (2008) since this is currently unsupported by experimental evidence), NorBC is the only nitric oxide reductase (19), and NosZ accepts electrons from menaquinol via complex III (25). Both thiosulfate and sulfide oxidation were modeled as complete oxidation to sulfate, transferring electrons to menaquinone instead of cytochrome c as previously described for the SOx complex (23). Neither polysulfide reduction nor formate oxidation were included as experimental evidence does not currently support these metabolic pathways, and reverse electron transport was modelled as discussed below.

Several postulated functions are not supported by direct biochemical evidence. Even for those protein complexes whose function has been identified in other organisms (e.g. cytochrome cbb_3_ oxidase), the stoichiometry of proton translocation *in vivo* is unknown for *S. denitrificans*. In these cases, several hypothetical mechanisms and/or proton pumping stoichiometries were considered. To identify which was most realistic, predicted relative growth efficiency (RGY; see Supplemental Methods for detailed derivation) was compared with yields of *S. denitrificans* grown in continuous culture (Table 2, (14)). If the predicted growth efficiency was close to the experimental data, the mechanism was considered plausible and vice-versa. We also inferred maximal chemosynthetic growth efficiency (CGE) by estimating the fraction of electrons required for carbon fixation via the reverse tricarboxylic acid (rTCA) cycle vs. other forms of metabolism (non-rTCA-PMF). A value for non-rTCA-PMF was obtained by comparing RGY between the hydrogen/nitrate condition and thiosulfate/nitrate condition in consideration of the overall metabolic model framework (Figure 1; see supplementary methods and Figs S1 & S2). Once this additional energy required for biosynthesis was determined, we could then derive a range of theoretical CGE values (Table 1).

For additional annotation of the *Sulfurimonas denitrificans* genome to support genome-based inferences, operons were predicted using OperonPredicter (65) and transmembrane domains were predicted using the TMHMM program (66). Further details for justifying choices made for the modeling are presented in the supplementary methods.

### Growth yield measurements in continuous culture

A custom-built chemostat was used to cultivate *S. denitrificans* (part numbers and design are available upon request). Triplicate temperature-controlled vessels with a liquid volume of 200 mL were fed via a peristaltic pump at an average dilution rate of 0.57 d^−1^ (the maximum growth rate of *S. denitrificans* is 1.44 d^−1^, (14)). Medium (see supplement) was kept anoxic by constant flushing with an anoxic gas mixture (N_2_/CO_2_ or H_2_/CO_2_) at a slight overpressure (27.5 KPa). Cultures were stirred constantly with teflon-coated magnetic stir bars, and all wettable surfaces were either glass, teflon, PEEK tubing (Idex Health & Science), or pharmed tubing (Cole-Parmer, USA). Light exposure was minimized by wrapping glass cultivation vessels with aluminum foil.

For cultivation with H_2_/NO_3_^−^, thiosulfate was omitted and H_2_/CO_2_ (80:20) was the headspace gas. Cultivation was carried out in a fume hood to prevent accumulation of flammable gas. An excess of dissolved hydrogen in the medium vessel was confirmed during cultivation by headspace extraction of a small volume of filtered culture and injection into a gas chromatograph. For cultivation with S_2_O_3_^2−^/NO_3_^−^, the headspace gas was N_2_/CO_2_ (80:20). For cultivation with S_2_O_3_^2−^/O_2_, a microaerobic environment was created by mixing N_2_/CO_2_ (80:20) with pure O_2_ to a final concentration of ~1% air saturation using a mass-flow controller (Tylan 260 series with RO-28 control box). Oxygen concentrations in chemostat vessels were monitored in real time with Pts3 optodes (Presens, Germany). The medium recipe and additional details are provided in the supplemental methods.

Cells fixed with borate-buffered formalin were stained with DAPI, filtered onto black 0.2 μM M polycarbonate filters, and counted with epifluorescence microscopy (67). Once numbers stabilized after 5-8 days of cultivation, both source medium and chemostat vessels were sampled for cell carbon (CC) and ash-free dry weight (AFDW; see SI for more detail).

For comparisons between hydrogen/nitrate and thiosulfate/nitrate, growth yield data were normalized to total nitrate consumed as determined by chemiluminescent detection using a NoxBox instrument (Teledyne, San Diego CA, USA) following the original protocol (68). For comparisons between thiosulfate/nitrate and thiosulfate/oxygen, yields were normalized to total sulfate produced as measured by ion chromatography with suppressed conductivity detection. *In silico* model results were compared to ratios of growth yields measured in continuous culture (this study, (14)) and overall growth yields (21).

## Acknowledgements

Funding was provided by NSF grants OCE-1136727, OCE-1559198, and the *WHOI Investment in Science Fund* (SMS) and fellowships awarded to JM from the National Sciences and Engineering Research Council of Canada (PGSM-405117-2011, PGSD-439487-2013) and the National Aeronautics and Space Administration Earth Systems Science Fellowship (PLANET14F-0075), an award from the Canadian Meteorological and Oceanographic Society, as well as funding from WHOI academic programs. We thank Ying Zhang for a critical reading of an earlier version of this manuscript and for advice on bioinformatics, François Thomas for invaluable help with qPCR, Florian Götz and Dali Smolsky for help with chemostat cultivation, Jeff Seewald and Sean Sylva for assistance with GC/IC measurements, Mak Saito for lending his lab’s mass-flow controller for microoxic cultivation, Carl Johnson for conducting the cell carbon measurements, and Scott Wankel and Carly Buchwald for assistance in making nitrate measurements. We especially thank Jörg Simon, Mårten Wikström, and Martin Klotz for helpful discussions regarding modelling and bioenergetics.

## Supplementary Information

**This PDF file includes:**

### Supplementary methods

**Table S1:** Hypotheses derived from the model and suggested tests for experimental validation.

**Table S2:** CGE values determined from chemostat cultivation.

**Figures S1 and S2:** Plots of predicted growth yield ratios (RGY) versus non-rTCA-PMF for H/ N:T/N and T/O:T/N ratios.

**Figures S3 and S4:** Expression of core metabolic genes in *S. denitrificans* under T/N and H/N conditions based on qPCR.

**Figure S5:** Sequence alignment of the *T. maritima* NfnB protein with Suden_1824 (the subunit that substitutes for NuoF in *S. denitrificans*).

**Figure S6:** Sequence alignment of both NuoG subunits in *S. denitrificans*, showing the missing regions in one of the copies.

**Figures S7, S8 and S9:** Growth curves of *S. denitrificans* under chemostat cultivation.

**Supplemental Dataset 1:** Descriptions of samples used for RNA extraction. **Supplemental Dataset 2:** Primers and targeted genes for qPCR analyses on batch cultures.

**SI references**

### Other supplementary material for this manuscript includes the following

1. qPCR sample and primer information excel sheet: “190619_supplemental_dataset_qPCR.xls”
2. Code for generating plots, raw sequences for alignments, as well as example calculations for CGE as a jupyter notebook can be found online here: https://osf.io/x8e5j

### Supplementary methods

#### Further information on model construction/inference

By only considering previously published mechanisms of sulfur oxidation, it was not possible to account for growth with thiosulfate and nitrate, despite being the original condition this organism was isolated under (1). This was because no electron transport pathway could link thiosulfate with the membrane quinone pool – necessary for the initial step of nitrate reduction. In the genome description of *S. denitrificans*, the SOx system was suggested to be responsible for thiosulfate oxidation (2). However, the SOx system is currently thought to be a soluble periplasmic membrane complex (3), and therefore cannot donate electrons to the quinone pool. Although a thiosulfate:quinone oxidoreductase responsible for partial oxidation of thiosulfate to tetrathionate has been described (4), neither this complex nor any other known thiosulfate-oxidizing enzymes are present in the genome of *S. denitrificans.* Therefore, we believe it is probable that a yet-undescribed complex serves to link the SOx system to the membrane quinone pool in *S. denitrificans*. Considering this, we employed a mechanism whereby the complete eight-electron oxidation of thiosulfate and sulfide to sulfate was linked to the reduction of menaquinone via the SOx system (Fig 1a).

To support carbon fixation, NAD(P)H and ferredoxin are both required for the reverse tricarboxylic acid (rTCA) cycle, yet no mechanism for ferredoxin production is known for *S. denitrificans* (5). However, the heterotrophic campylobacterium *Campylobacter jejuni* possesses a non-canonical NADH:ubiquinone oxidoreductase complex (complex I) that oxidizes reduced flavodoxin instead of NADH (6). Such a complex has the potential to reduce ferredoxin if run in reverse; indeed, *S. denitrificans* has a non-canonical complex I operon similar to *C. jejuni.* However, NAD(P)H is also needed, and not produced by such a mechanism. Therefore, for modeling purposes, we included a hypothetical mechanism that reduces both NAD(P)+ and ferredoxin simultaneously via an electron-bifurcation-like mechanism (Fig 1d). Further details on this mechanism are discussed in the main text.

#### Energy Conservation Efficiency of Enzymatic Complexes

We then considered growth under conditions previously described for *S. denitrificans* (2, 7, 8). Growth yields were estimated from combining core dissimilatory reactions, reversed electron transport, and an overall reaction for the rTCA cycle (Table 1). However, using canonical proton-pumping stoichiometries for these complexes resulted in several discrepancies. Firstly, growth efficiencies did not differ between thiosulfate and hydrogen oxidation, in contrast to data previously reported in batch culture (8) and chemostat growth data presented here (Table 2). Secondly, the *in silico* ratio between aerobic and denitrifying growth efficiency was much higher than what has previously been determined in chemostat experiments (7).

The first discrepancy – why modeled growth yields *in silico* were identical during sulfur and hydrogen oxidation but different in growth experiments - was resolved by considering the inferred structure of enzyme complexes. The hydrogenase in *S. denitrificans* likely has a transmembrane domain - allowing it to accept protons from the periplasm during quinone reduction (Fig 1b) (2). When coupled to a quinol-accepting reductase, this translocates protons via a redox-loop mechanism (9). In contrast, proteins catalyzing sulfur oxidation coupled to quinone reduction have not been demonstrated to translocate protons. This is because they either do not have transmembrane domains (10) or the quinone-accepting site and the redox module both face the cytoplasm and thus do not translocate protons (4, 9). Since thiosulfate oxidation likely occurs in the periplasm in *S. denitrificans*, we chose to model the SOx system with a periplasmic-facing non-transmembrane anchor similar to sulfide:quinone oxidoreductase (SQR; Fig. 1a). With this mechanism in place, the ratio between hydrogen oxidation and sulfur oxidation under denitrifying conditions was congruent with experimental data (Tables 1 and 2).

The second inconsistency – large differences in growth yield ratios between aerobic and denitrifying respiration, but only a small difference observed during experiments - could be reconciled if the energy conservation efficiency of aerobic respiration was either lower or that of denitrification was higher. Although it was previously thought that proton translocation is not catalyzed by denitrification enzymes (11), recent studies have suggested that denitrification is net electrogenic through the concerted action of qcrABC (Complex III) and denitrification enzymes (12, 13). However, since this net reaction for denitrification is thought to have a H+/e− of 1, it would result in a much higher RGY ratio than observed for *S. denitrificans* if aerobic respiration occurred with the same efficiency as for typical aerobes (H+/e− = 3; (9)).

Therefore, the efficiency of aerobic respiration coupled to sulfur oxidation must be lower to account for these observations. If the SOx system directly donated electrons to soluble cytochrome c under aerobic conditions (14), this would reduce overall efficiency by bypassing the energy conserving complex III. However, this would require that complex III operates in both the forward and reverse direction. This occurs in the autotroph *Acidithiobacillus ferrooxidans,* but it possesses one dedicated complex for each direction (15). Given that *S. denitrificans* has only one copy of complex III (2), we rejected this possibility in the model.

The only plausible explanation remaining is that the complex predicted to catalyze aerobic respiration in *S. denitrificans* (a cbb_3_ high-affinity type cytochrome oxidase; (2)), has reduced bioenergetic efficiency. While this complex is theoretically capable of proton transport, it has been shown to be unable to pump protons effectively against membrane potentials typically encountered in living cells (16–20). Therefore, under physiological conditions, it appears likely that *S. denitrificans*’ cbb_3_ cytochrome oxidase does not pump protons.

Even without pumping protons, the cbb_3_ complex would still conserve more energy than denitrification enzymes because protons for oxygen reduction typically come from the cytoplasm, contributing to the proton-motive force and a H+/e− of 1 (21). When coupled with Complex III, aerobic respiration would have an overall H+/e− of 2, which would lead to predicted RGY yields that are consistent with experimental observations (Table 2). An alternative (but in our opinion less likely) explanation for these observations is that “chemical” protons for oxygen reduction are abstracted from the periplasm. This has not yet been demonstrated for any cytochrome c oxidase, but it is known that NO reductases obtain protons for NO reduction from the periplasm, and that they may be the evolutionary precursor of complex IV (22, 23). If this is true, it follows that there may be some extant cytochrome c oxidases that obtain their “chemical” protons from the periplasm. Indeed, such a discovery would serve as a “missing link” between NO and O_2_ reduction and reinforce this evolutionary scenario (Mårten Wikström, pers. comm.). Only biochemical experiments that specifically investigate these mechanisms can resolve this uncertainty (Table S1). With these hypothetical mechanisms in place (Fig 1), the model reproduced experimental growth yield data satisfactorily, and was considered to be complete.

#### Calculation of theoretical chemosynthetic growth efficiency (CGE)

We also calculated CGE, which is a parameter that can be used for modelling chemosynthetic primary productivity in natural environments (24, 25). CGE is the fraction of electrons that go to assimilatory carbon fixation as a proportion of the total electrons oxidized. To do so, we exploited a unique property of our conceptual model that allowed for the calculation of the proton-motive force needed for cell growth, maintenance, motility etc., in addition to that needed for carbon fixation by the rTCA cycle (hereafter “non-rTCA-PMF”). To estimate non-rTCA-PMF, we first assumed that the ratio of non-rTCA-PMF to carbon fixed is consistent between different combinations of electron donors and acceptors. In other words, we are assuming that cell composition and biosynthetic reactions are constant between different growth conditions. Considering the equations in Table 1, we further assume that to fix one molecule of oxaloacetate from CO_2_, *S. denitrificans* needs to translocate 25 protons to account for ATP and reductant required.

We then compared the RGY between T/N and H/N conditions to determine CGE. We chose not to use the T/O : T/N ratio because of uncertainties with regards to the potential negative effects of oxygen that may reduce CGE (25). The ratio of potential growth yields between H/N and T/N conditions will vary as a function of the amount of non-rTCA-PMF needed (Fig S1). The maximum ratio will be obtained where non-rTCA-PMF is zero. This is because for growth under T/N conditions, our model predicts that the process of transferring electrons from thiosulfate to menaquinone and to complex I is electroneutral. In contrast, under the H/N condition, electrons derived from H_2_ oxidation can generate a proton motive force by charge separation during reverse electron transport ((9); Fig 1d). This does not mean that reverse electron transport is net electrogenic under H/N conditions, but rather that the process of obtaining reductant for carbon fixation under H/N conditions does result in the translocation of protons, which can be used for reversed electron transport. Because of this, carbon fixation is much more efficient for the H/N condition relative to T/N, leading to a maximum theoretical ratio of 3.33 (i.e. 7.5 electrons are needed to produce one molecule of oxaloacetate from CO_2_ under the H/N condition, whereas 25 are needed for the T/N condition). However, the actual value will always be below this value since the cell requires PMF/ATP for other biosynthetic reactions (what we term “non-rTCA-PMF”), which we predict will occur at a fixed efficiency (H+/e− = 2; Fig 1b).

In our experiments, the measured Y_H/N_:Y_T/N_ ratio was between 2.3-2.4. According to the conceptual model described above, this ratio can be written as follows (where the product of x and its coefficient represents the amount of “non-rTCA-PMF” needed, with a coefficient added to take into consideration the different proton-pumping efficiencies of the H/N and T/N conditions):

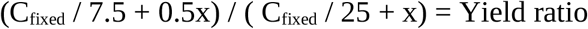

which simplifies to:

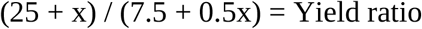

Solving for x by substituting both RGY values we measured (Table 2), we can calculate f(2.3) = 35, f(2.4) = 52. This allows us to calculate theoretical CGE values based on the production of one molecule of oxaloacetate from CO_2_ (requiring 10 electrons). An example is shown below for T/N conditions:

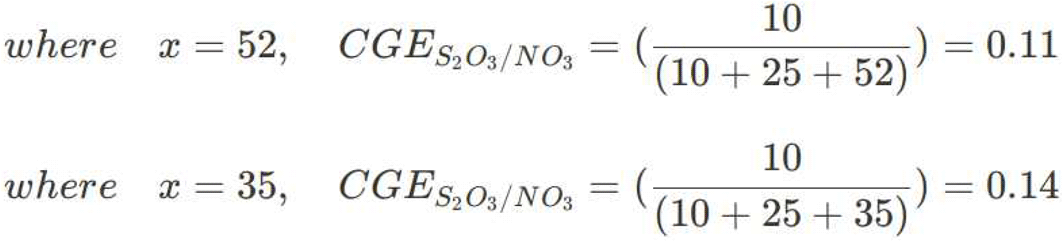

CGE_T/N_ = 0.11(10/(10+25+52)) − 0.14 (10/(10+25+35)). The lowest value is slightly higher than Klatt and Polerecky’s estimates for *S. denitrificans* based on the data reported in (7) (0.098), but close to those estimated by the same authors for other *Sulfurimonas* cultures (0.148; 0.125; 0.113; (24)).

CGE_T/O_ = 0.20 (10 / (10+12.5+26)) − 0.25 (10 / (10 + 12.5 + 17.5). These values are slightly higher than the Klatt and Polerecky value for *S. denitrificans* calculated from the data reported in (7) (0.167), but this difference could perhaps be attributed to oxidative damage caused by O_2_ ((25) and references therein). No other CGE measurements are available for *Sulfurimonas* strains under microaerobic conditions (since our CGE measurements are likely underestimates; see discussion below).

CGE_H/N_ = 0.23 (10 / (10 + 7.5 + 26)) − 0.28 (10 / (10 + 7.5 + 17.5)). No current data is available for cultivars for CGE, but these values are congruent with highest CGE value reported for our previously published incubations of natural microbial communities dominated by *Sulfurimonas* (0.27, for incubations with H_2_ and low oxygen; (25)).

CGE_H/O_ = 0.31 (10 / (10 + 5 + 17.33)) − 0.375 (10 / (10 + 5 + 11.66)). No current data is available for cultivars growing with hydrogen and oxygen, and is complicated by the negative effect that oxygen has on these organisms, meaning that optimal cultivation under this condition would require a more sophisticated experimental setup that could provide oxygen at low, but non-limiting concentrations.

We were unfortunately unable to compare these theoretical estimates to our chemostat data since our actual measurements of CGE were lower than previously reported (Table S2). We believe this is likely to be an experimental artefact that underestimated actual CGE. Nitrate limitation was not achieved at steady-state in our continuous culture experiments, despite an excess of thiosulfate/hydrogen. It is possible that constant flushing with gas during chemostat growth removed trace amounts of oxygen needed for biosynthesis (26) or some other limiting nutrient and accounted for lower CGE values we observed (approximately 1/2 of expected according to our model and previous observations). Other potential explanations for low CGE include the filters used to capture biomass (nominal pore size of 0.3 μm) not retaining all cells, excretion of fixed organic matter into the media. The latter two possibilities seem unlikely given that Timmer-Ten Hoor (1981) used filters with larger pore sizes and excretion would be expected to also occur in their experiments. Alternative explanations include non-optimal nutrient stoichiometry in the growth media, or the introduction of a toxic/nutrient-chelating substance into the medium from the gas or cultivation vessels/tubing. Another possibility is that differences in dilution rate affects yield. Little is known about this question, but the available data (Fig 1 in (7)) suggest that absolute yield differences should be minor (~5%) between the dilution rate employed in our study and in (7). Despite these limitations, our values for RGY are nearly identical to those previously reported and consistent with the model.

#### Chemostat cultivation

The chemostat was operated as described in the main text. The medium was made by first dissolving and autoclaving the following components in MilliQ water (per L final volume): NaCl 10 g; NH_4_Cl 1 g, MgSO_4_∙7H_2_O 3.5 g; CaCl_2_∙2H_2_O 0.42 g; KCl 0.7 g; KNO_3_ 2.0 g; Trace element solution SL-6 (DSMZ 465) 1mL. After autoclaving, separately-sterilized anaerobic solutions of the following components were added (per L final volume): NaHCO_3_ 5 g; KH_2_PO_4_ 0.5 g; selenate/tungstate solution (DSMZ 944a) 4 mL; 0.7 mL of 2 mg∙mL^−1^ FeCl_3_ in 0.1 NH_2_SO_4_; Na_2_S_2_O_3_∙5H_2_O 5 g. The thiosulfate solution was sterilized by filtering through a 0.2 μm membrane and the bicarbonate solution was autoclaved under a N_2_/CO_2_ (80:20) atmosphere.

For all incubation conditions, pH measured daily was 7.19 ± 0.04 for H_2_/NO_3_^−^, 7.06 ± 0.01 for S_2_O_3_^2−^/NO_3_^−^ and 7.02 ± 0.02 for S_2_O_3_^2−^/O_2_. Upon sampling, cell numbers had not varied on average by more than 11 % over at least 2 days of cultivation, indicating steady state had been reached (Fig. S7, S8 and S9). This occurred after 7, 8, and 5 days for H_2_/NO_3_^−^, S_2_O_3_^2−^/NO_3_^−^, and S_2_O_3_^2−^/O_2_ respectively.

Yield measurements (cell carbon, CC; ash-free dry-weight, AFDW) were conducted as follows. Combusted and pre-weighed GF-75 filters (0.3 μM M pore size; Advantec) were used for both measurements. For AFDW, filters were first wetted with 0.5 M ammonium formate (27). Next, 20 mL of medium or cells were filtered, followed by 10 mL of ammonium formate. Cells were dried overnight at 80°C, placed in a vacuum desiccator, and weighed when cool. Then, filters were combusted at 480°C for 8 h and weighed as before. Medium blanks were subtracted to determine AFDW. For cell carbon, 20 mL of both medium and cell culture were filtered using pre-combusted glassware. Filters were stored in pre-combusted shell vials in −80°C prior to analysis. Total cell carbon was determined by acidification and subsequent combustion of biomass on a Carlo Erba/Fisons 1107 Elemental Analyzer as previously described (25).

#### Batch Cultivation for qPCR Experiments

Media was autoclaved for 20 min in a Widdel flask, cooled under anoxic N_2_ gas, and transferred to N_2_-purged Hungate tubes, serum vials or nalgene flasks and sealed with butyl stoppers. pH was adjusted to 7.0+/−0.05 before autoclaving by adding 3mL of a solution of 1N NaOH. Bicarbonate and thiosulfate solutions were prepared in anoxic N_2_-purged water and added after autoclaving through a sterile 0.2 μm filter. In addition, 0.6 mL of a sterile 0.1N H_2_SO_4_ solution containing 2mg FeCl_3_ × 5 H_2_O was added after autoclaving, and 1mL of trace metal solution SL-6 was added prior to autoclaving.

For hydrogen-grown cultures, thiosulfate was omitted from the medium, and the headspace was evacuated using a vacuum pump and refilled with H_2_/CO_2_ (80/20%) to 28psi gauge pressure. Thiosulfate-grown cultures were filled with 5psi gauge pressure N_2_/CO_2_ (80/20%). A solution of anoxic sodium bicarbonate was used to buffer the pH at 7 where CO_2_ was used in the headspace. For each condition, quadruplicate cultures were grown, with an additional uninoculated control to account for abiotic processes. Pressure was measured periodically in vessels using a pressure gauge fitted to a luer-lok fitting to which a sterile needle was attached. pH was monitored by taking subsamples using syringes and measuring using a desktop probe.

Cell growth was monitored by direct counts as described above. Cells were harvested in both exponential and stationary phase, and 1 sample from each phase was taken from each treatment (S_2_O_3_^2−^/NO_3_^−^ and H_2_/NO_3_^−^) for RNA extraction. In addition, the S_2_O_3_^2−^/NO_3_^−^ grown cells used to inoculate the H_2_/NO_3_^−^ culture were sampled to observe changes in gene expression associated with adaptation to H_2_ as an electron donor.

#### Culture Quality Control

The presence of heterotrophic contamination was tested by incubating cells in anaerobic media containing nitrate, 0.5% tryptone, 0.5% glucose and 0.5% yeast extract. Growth was also assayed with formate (20mM) as sole electron donor with nitrate (20mM) as electron acceptor.

To test for contaminants that could not be detected by the method above, DNA was extracted from 4 samples (1 frozen culture, and 3 growing, replicate cultures) using the UltraClean Microbial DNA Isolation Kit (Mobio, Carlsbad, CA). 16S DNA was amplified using the 341F/907RC primer pair (Schäfer *et al.*, 2000) with a GC-clamp designed for DGGE using Fermentas Taq polymerase (Fisher Fermentas, Vilnius, Lithuania) and the following conditions: (94°C for 5 min, 20 cycles of 94°C 1min, 65°C 1min (touchdown 0.5°C/cycle) and 72°C 1min, followed by 15 cycles of 94°C 1min, 55°C 1min and 72°C 1min, followed by 7 min at 72°C). The PCR product was cleaned up using the Axyprep PCR Clean-up kit (Axygen, Union City, CA). 300ng of DNA from each sample was loaded onto a polyacrylamide gel as previously described (28) and stained using SYBR-safe DNA stain and visualized under UV light.

#### RNA preservation, extraction and reverse transcription

Quadruplicate samples taken during the growth experiments were preserved using RNAprotect bacteria reagent (Qiagen, Hilden, Germany) as per manufacturer instructions, and frozen at −80°C until RNA extraction, which was carried out within 2 weeks of preservation. RNA was extracted from triplicate samples (one replicate was not used) using the RNeasy mini kit (Qiagen). Yield was measured using a Nanodrop instrument (Fisher Scientific, Waltham, MA) and integrity was verified using gel electrophoresis. Residual DNA was digested using the Turbo DNAase kit (Ambion, Austin, TX) at 35°C for 1hr, keeping the RNA concentration constant between all digestions. RNA was then purified with the RNeasy Minielute RNA cleanup kit (Qiagen) and re-quantified using a Nanodrop. The absence of DNA contamination of the RNA was verified by conducting a PCR assay on the RNA using Fermentas Taq polymerase (Fisher Fermentas, Vilnius, Lithuania) and the following conditions: 95°C:2min, 35 cycles of 95°C:30sec, 55°C:40sec, 72°C:30sec followed by a 5 minute hold at 72°C. A no-RT control was also tested for amplification from two randomly selected RNA samples. Reverse transcription was accomplished using the DyNAmo cDNA synthesis kit (Thermo Fisher, Waltham MA) and the following thermocycling conditions: 25°C:10min, 37°C:60min, 85°C:5min. The amount of RNA added to each reaction was 134.4 ng. cDNA was diluted 20-fold before qPCR.

#### qPCR primer design and assays

PerlPrimer (29) was used to design primers for each specific gene (Supplemental Dataset 2) according to the parameters described in (30). In-silico specificity of each primer pair was tested by conducting BLAST searches against the *S. denitrificans* DSM 1251 genome. In-vitro specificity was tested by conducting a PCR amplification on genomic DNA and gel electrophoresis. Single bands were observed in all cases, indicating specific amplification. The linearity of response was tested by creating a standard curve using *S. denitrificans* genomic DNA (10^6^ copies to 10 copies per well). Efficiency was between 85 and 103.6% for all primer pairs. Genomic DNA was isolated from pure cultures of *S. denitrificans* using the UltraClean Microbial DNA Isolation Kit (Mobio, Carlsbad, CA).

qPCR assays were carried out on the Stratagene MX3005P instrument (Agilent, Santa Clara CA) using Fermentas Maxima qPCR Master Mix (Fisher Fermentas, Vilnius, Lithuania) with the following mixture per well: 7.5 μL master mix, 1.5 μL template, 1.5 μL primer mix (5 μM) and 4.5 μL H_2_O. Thermocycling conditions were as follows: 95°C:10min, followed by 40 cycles of 95°C:15s, 60°C:30s, 72°C:30s, then by 55°C:30s and a slow ramp to 95°C to construct disassociation curves. Each sample was run in duplicate, and standard curves were re-run with every plate. DNA amplification was quantified using SYBR green with ROX as a reference dye.

#### qPCR quality control and normalization

After thermocycling, wells were manually inspected for evaporation, and those that differed significantly from their duplicate were removed from further analysis. Amplification plots were manually inspected and baselines changed if necessary, and the critical cycle C_t_ was adjusted to make sure it was within the log-linear portion of the amplification plot. Absolute expression values were plotted with ggplot2 (code and raw data available at https://osf.io/x8e5j/).

#### Ion Chromatography

Culture samples were centrifuged at 10,000g for 2 minutes to pellet cells, and the supernatant decanted, and stored frozen at −20°C until analysis. Prior to analysis, samples were diluted 100-fold in Milli-Q water using a 4-place balance, and for the H/N samples, shaken overnight at 35°C to resolubilize any precipitate formed. Standards were prepared from commercially purchased standard solutions to bracket expected concentrations (1000ppm Specpure standards for NO_2_^−^, NO_3_^−^, Cl^−^, SO_4_^2−^: Alfa Aesar; 11200ppm S_2_O_3_^2−^: Fisher Scientific).

Samples were diluted 100-fold, and run on a Dionex DX 500 Ion Chromatograph equipped with a GP50 gradient pump, ED40 electrochemical detector, LC30 chromatography oven and a AS40 autosampler. Each chromatogram was manually adjusted to ensure peaks were integrated correctly.

Quantification of anions was accomplished by comparing the integrated peak size with a standard curve generated from the above standards.

**Table S1.** Hypothetical biochemical mechanisms and experimental tests

| Hypothesis | Predictions of hypothesis | Experimental test and expected result |
| --- | --- | --- |
| **SOx complex is attached to membrane and transfers electrons to quinone pool** | QcrABC (complex III) solely responsible for energy conservation during sulfur oxidation; in contrast, hydrogen oxidation can conserve energy via redox loop mechanism | **Knock out or inhibit complex III: Thiosulfate-nitrate dependent growth should stop; hydrogen-nitrate dependent growth may continue with lower yield if NapGH can act as a quinol:acceptor reductase independently (Kern and Simon 2008), but denitrification will not proceed beyond nitrite** |
| **cbb<sub>3</sub> cytochrome c oxidase does not pump protons under physiological conditions\*** | Proton-motive force during oxic respiration comes from the combined action of QcrABC and the abstraction of protons from the cytoplasm by cbb<sub>3</sub> | **Uncouplers of cbb<sub>3</sub> proton pumping should not affect physiological growth yields** |
| **NADH:ubiquinone oxidoreductase is responsible for NAD(P)<sup>+</sup> and ferredoxin reduction** | Complex donates electrons to both NAD(P)<sup>+</sup> and ferredoxin during reverse electron transport | **Use inverted vesicles with complex I and ATPase: Reduction of ferredoxin and NAD(P)<sup>+</sup> should be possible with ATP present to generate pmf** |

\*An alternative possibility is that “chemical protons” for cbb<sub>3</sub> are abstracted from the periplasm, which would result in the same growth yields.

**Table S2.**
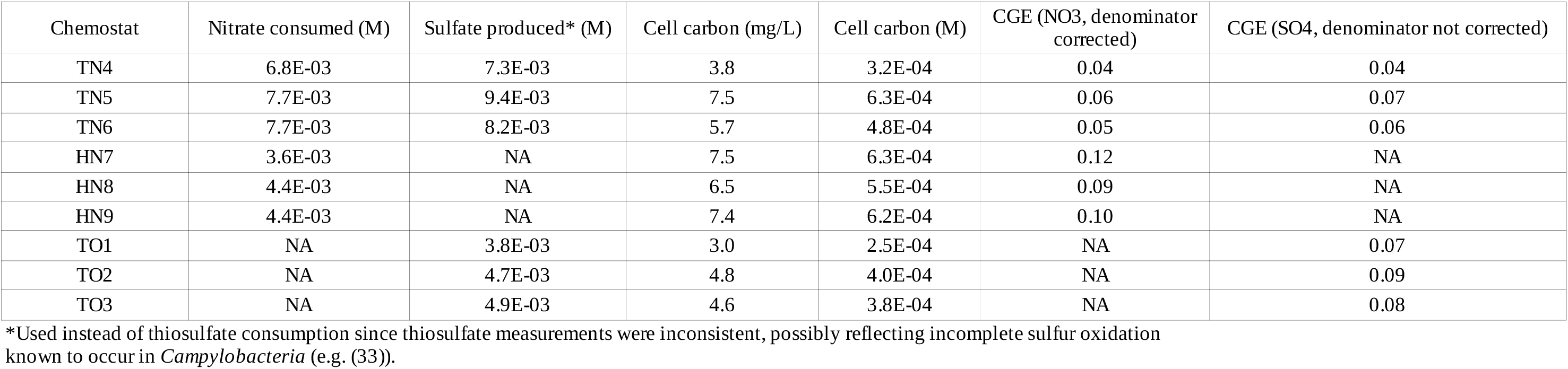
Calculations of Chemosynthetic Growth Efficiency from chemostat growth experiments. TN=Thiosulfate/nitrate, HN=Hydrogen/nitrate, TO=Thiosulfate/oxygen.

| Chemostat | Nitrate consumed (M) | Sulfate produced* (M) | Cell carbon (mg/L) | Cell carbon (M) | CGE (NO3, denominator corrected) | CGE (SO4, denominator not corrected) |
| --- | --- | --- | --- | --- | --- | --- |
| TN4 | 6.8E-03 | 7.3E-03 | 3.8 | 3.2E-04 | 0.04 | 0.04 |
| TN5 | 7.7E-03 | 9.4E-03 | 7.5 | 6.3E-04 | 0.06 | 0.07 |
| TN6 | 7.7E-03 | 8.2E-03 | 5.7 | 4.8E-04 | 0.05 | 0.06 |
| HN7 | 3.6E-03 | NA | 7.5 | 6.3E-04 | 0.12 | NA |
| HN8 | 4.4E-03 | NA | 6.5 | 5.5E-04 | 0.09 | NA |
| HN9 | 4.4E-03 | NA | 7.4 | 6.2E-04 | 0.10 | NA |
| TO1 | NA | 3.8E-03 | 3.0 | 2.5E-04 | NA | 0.07 |
| TO2 | NA | 4.7E-03 | 4.8 | 4.0E-04 | NA | 0.09 |
| TO3 | NA | 4.9E-03 | 4.6 | 3.8E-04 | NA | 0.08 |

\*Used instead of thiosulfate consumption since thiosulfate measurements were inconsistent, possibly reflecting incomplete sulfur oxidation known to occur in *Campylobacteria* (e.g. (33)).

**Figure S1:**
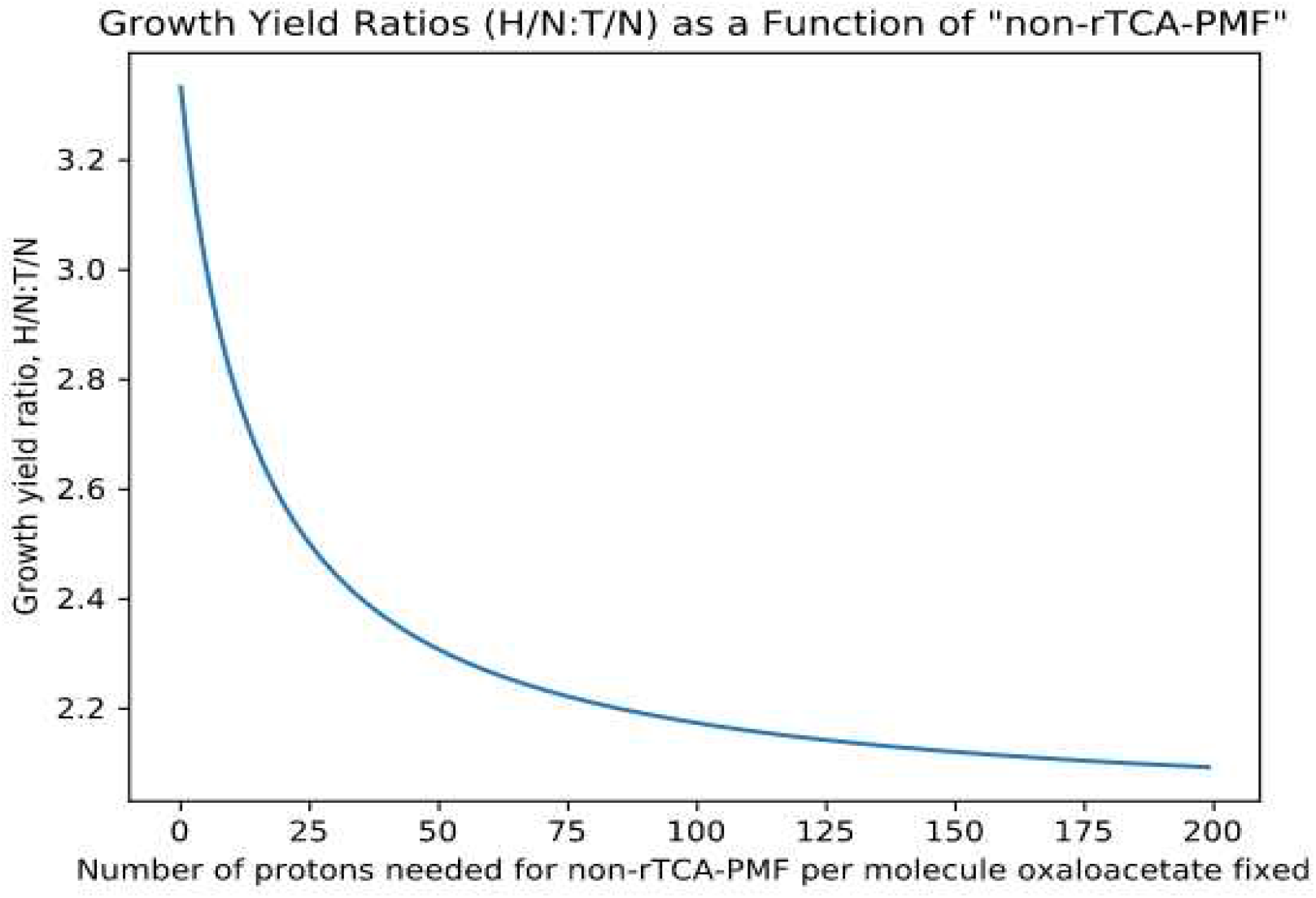
Predicted relative growth yields between H/N and T/N conditions depending on the fraction of non-rTCA-PMF required per molecule of oxaloacetate fixed. See jupyter notebook at https://osf.io/x8e5j for a more detailed explanation.

**Figure S2:**
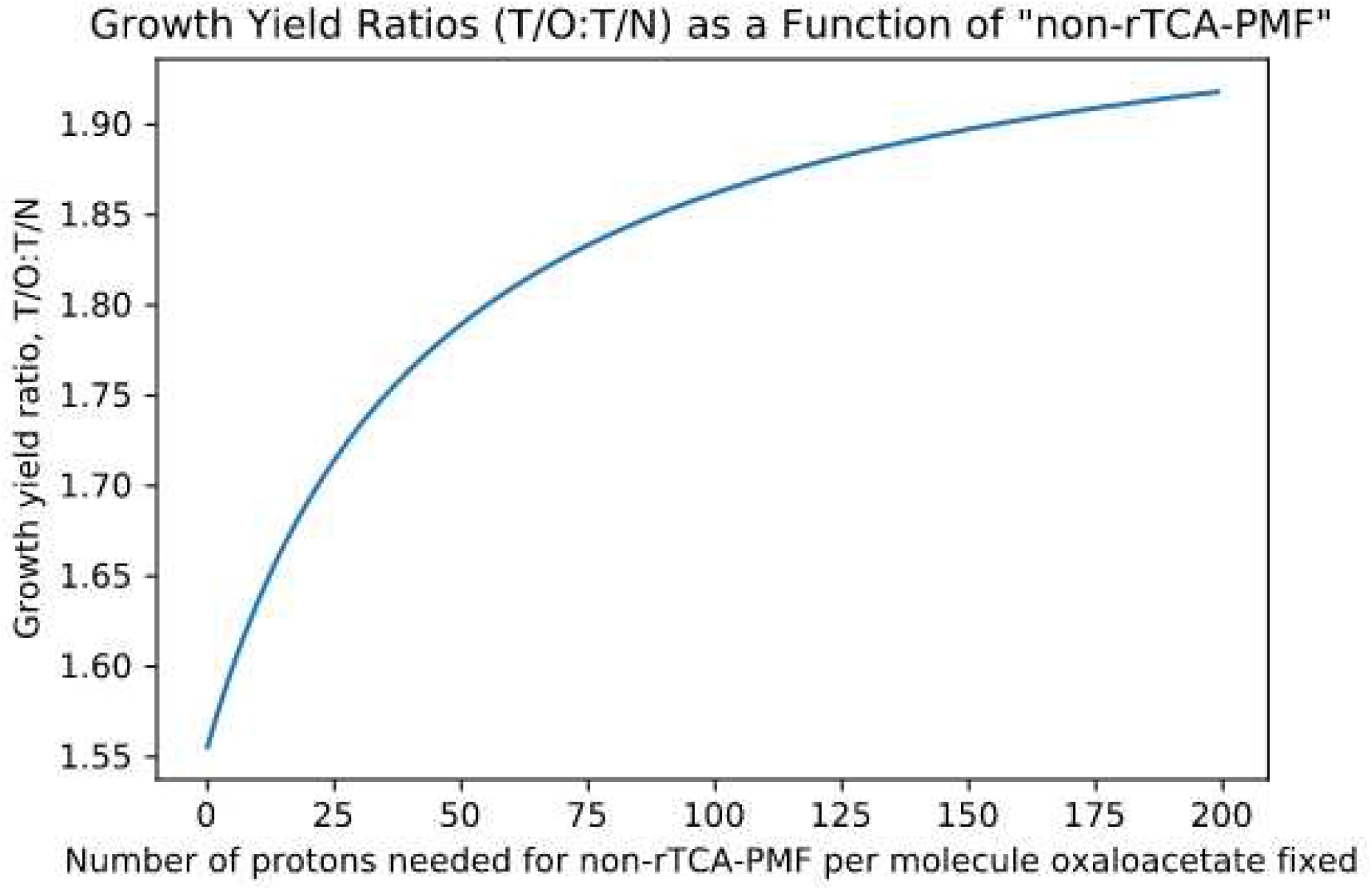
Predicted relative growth yields between T/O and T/N conditions depending on the fraction of non-rTCA-PMF required per molecule of oxaloacetate fixed. See jupyter notebook at https://osf.io/x8e5j for a more detailed explanation.

**Figure S3:**
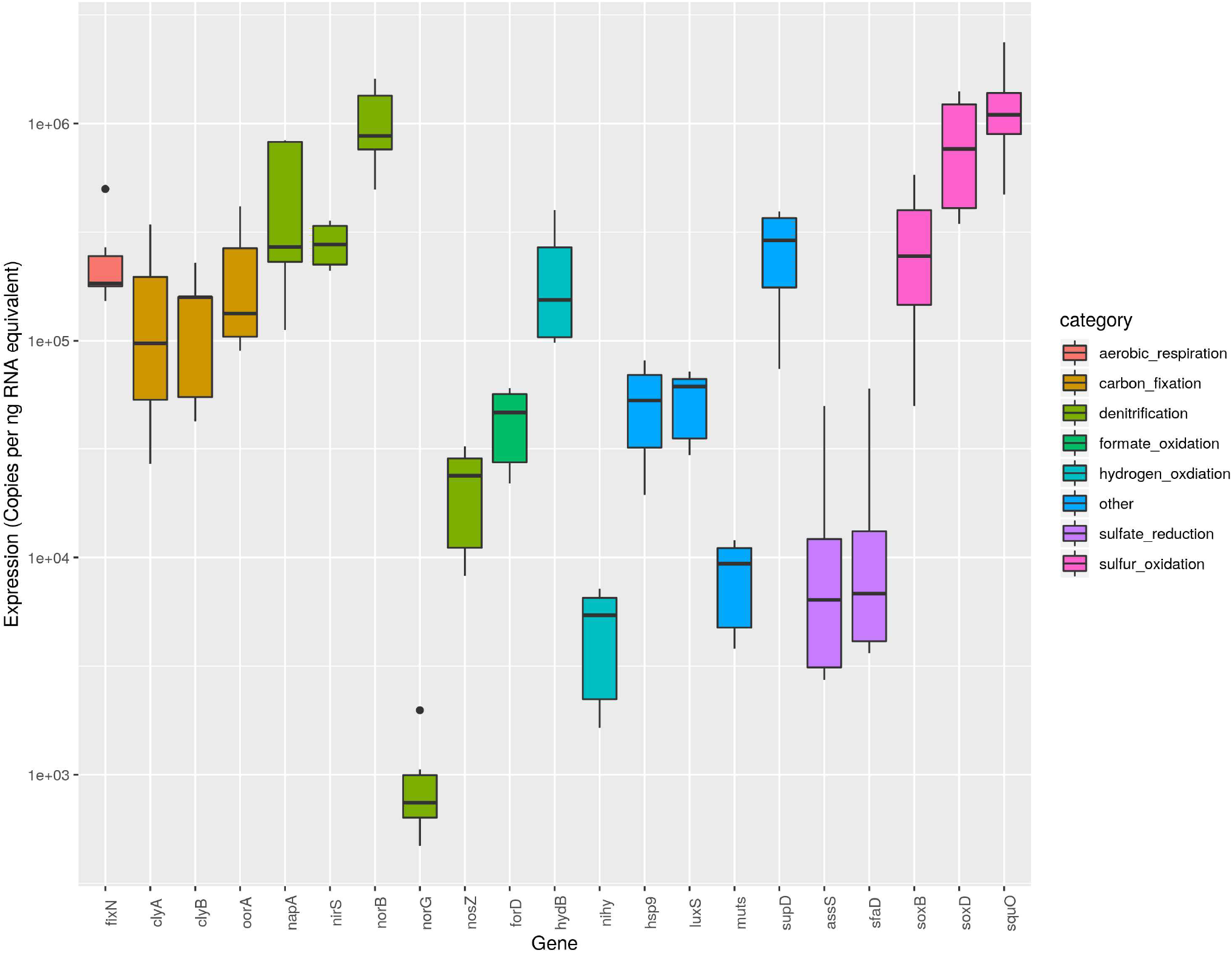
mRNA Expression under Thiosulfate/Nitrate Growth Condition (n=6) RT-qPCR transcript abundance of selected genes in *S. denitrificans*. Values for absolute expression are derived from 6 independent measurements from both thiosulfate/nitrate and hydrogen/nitrate-grown cells at two separate time points (exponential and stationary phase) and normalized to total RNA (see Supplemental Dataset 1 for more detail). Genes are colored according to their predicted metabolic role as shown in the legend. Gene abbreviations correspond to the genes identified in Supplemental Dataset 1. The sulfide:quinone oxidoreductase mentioned in the main text corresponds to “squO” (the last gene in the boxplot). See jupyter notebook at https://osf.io/x8e5j for plotting code.

**Figure S4:**
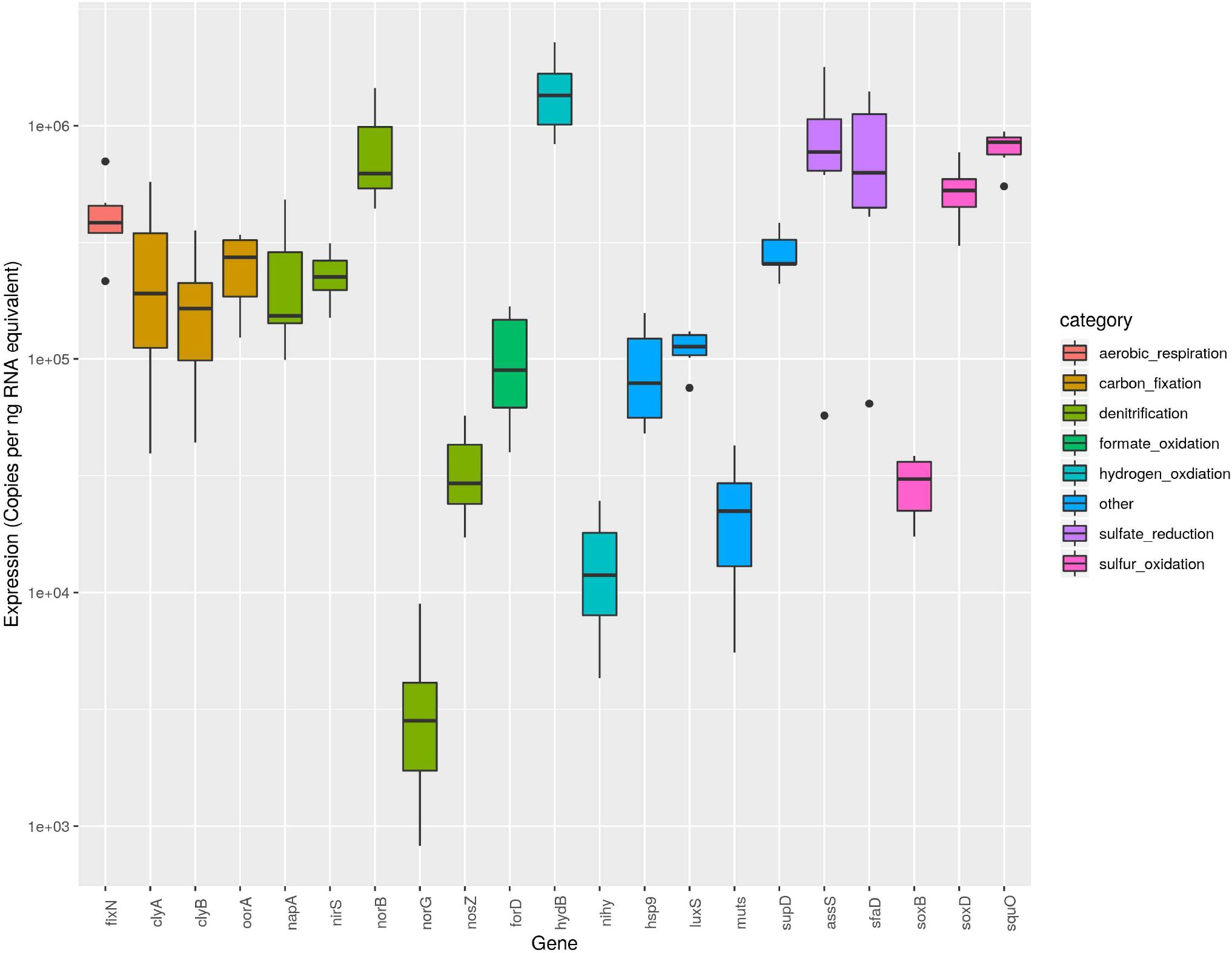
mRNA Expression under Thiosulfate/Nitrate Growth Condition (n=6) RT-qPCR transcript abundance of selected genes in *S. denitrificans*. Values for absolute expression are derived from 6 independent measurements from both thiosulfate/nitrate and hydrogen/nitrate-grown cells at two separate time points (exponential and stationary phase) and normalized to total RNA (see Supplemental Dataset 1 for more detail). Genes are colored according to their predicted metabolic role as shown in the legend. Gene abbreviations correspond to the genes identified in Supplemental Dataset 1. The sulfide:quinone oxidoreductase mentioned in the main text corresponds to “squO” (the last gene in the boxplot). See jupyter notebook at https://osf.io/x8e5j for plotting code.

**Figure S5:**
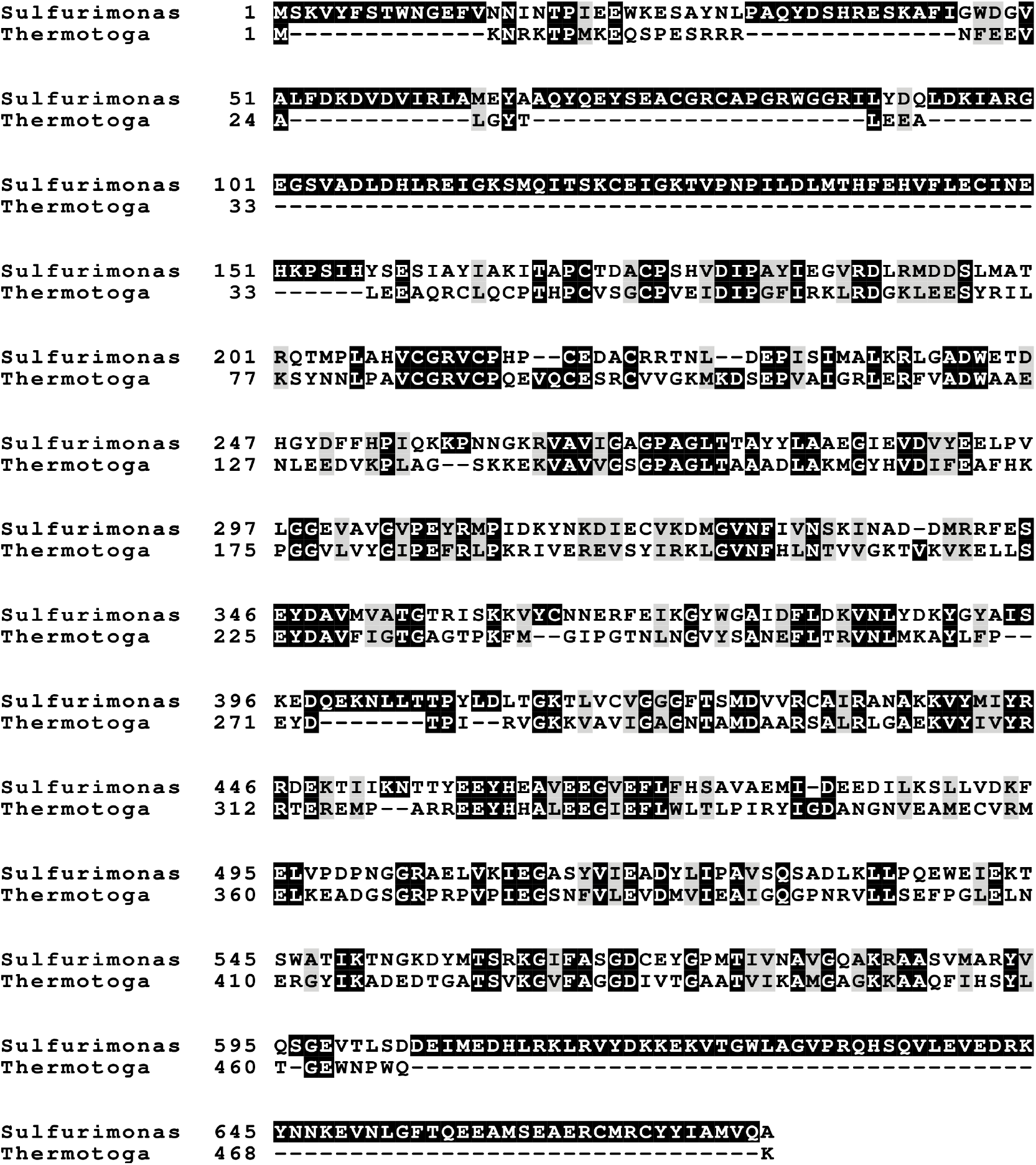
Sequence alignment of the *T. maritima* NfnB protein with Suden_1824 (the subunit that substitutes for NuoF in *S. denitrificans*). As mentioned in the main text, relevant residues for binding the phosphate group of NADP(H) (Tyr310, Arg311-312 in *T. maritima*) and cysteines ligating the iron-sulfur clusters that putatively pass electrons to ferredoxin are conserved in Suden_1824, as well as most residues involved in binding FAD (31). See https://osf.io/x8e5j for raw alignments.

**Figure S6:**
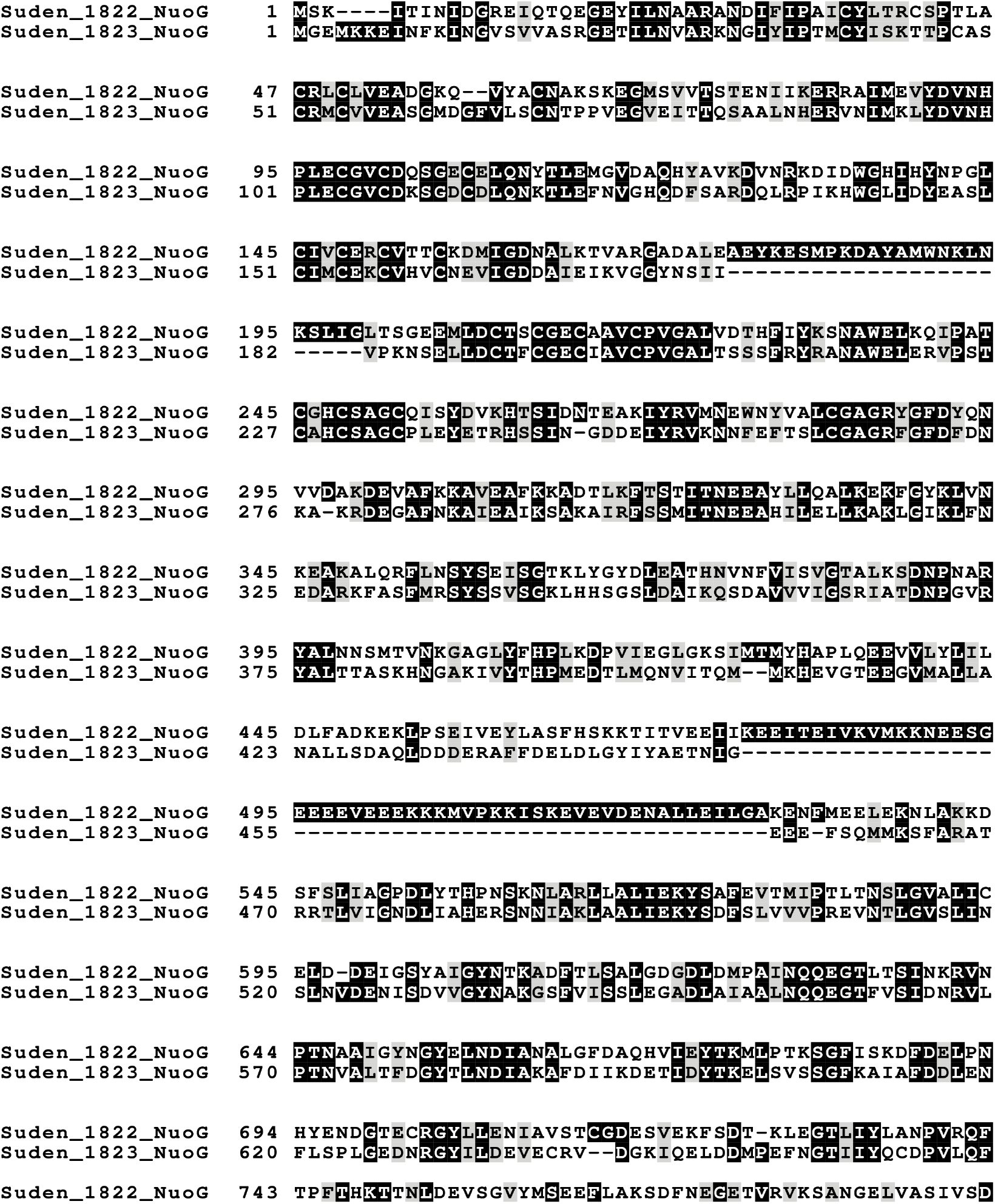
Sequence alignment of both NuoG subunits in *S. denitrificans,* showing the missing regions in one of the copies. As noted in the main text, the shorter subunit (Suden_1823) is “missing” two main regions. The first (AA 466-532 of Suden_1822) contains no obvious functional motifs. The second missing region (AA 175-204 of Suden_1822) is found between two cysteine residues that putatively bind an additional iron-sulfur cluster (32). See https://osf.io/x8e5j for raw alignments.

**Figure S7:**
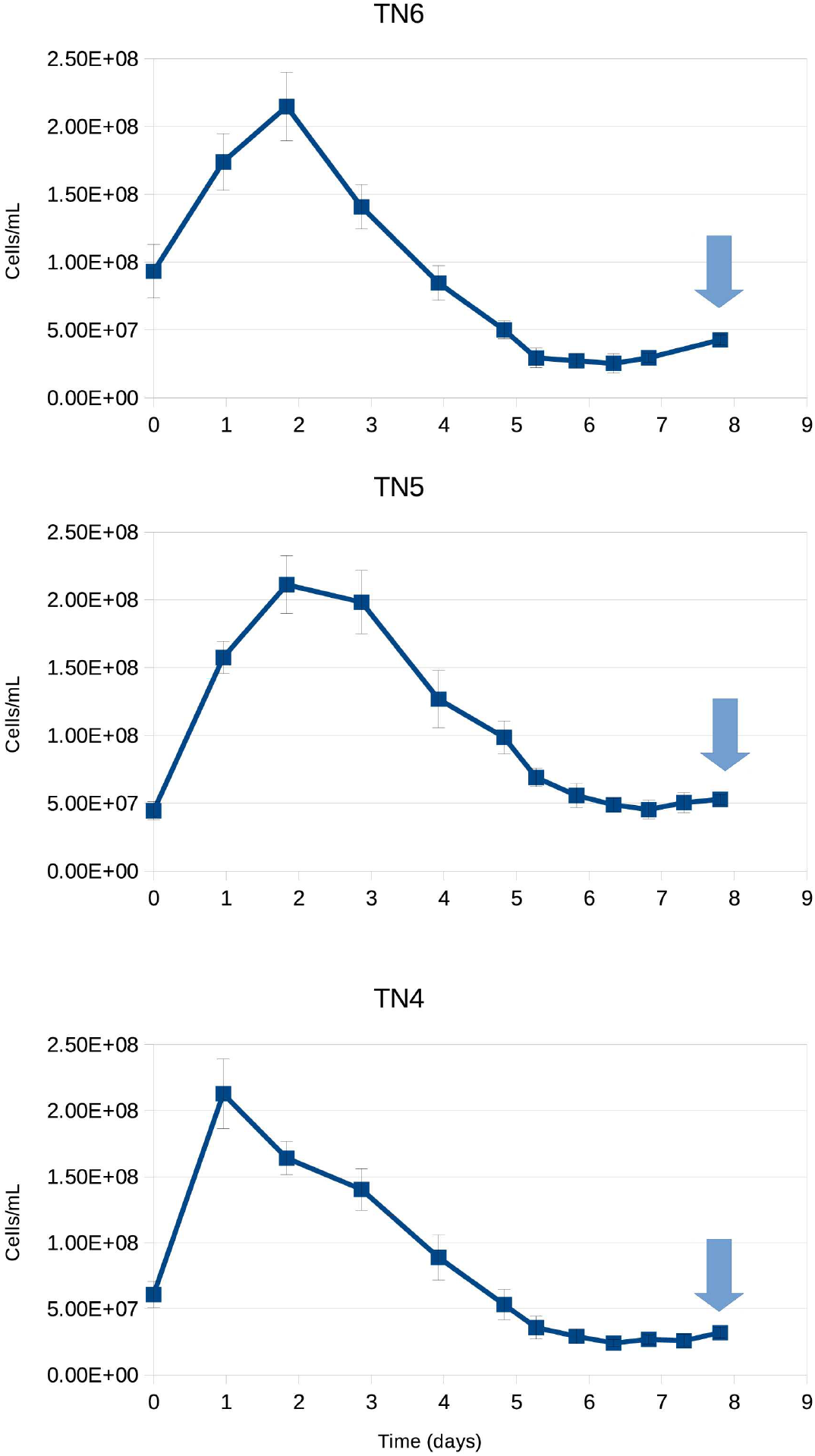
Growth curves for three replicate chemostat experiments grown with thiosulfate and nitrate. Arrows indicate time of sampling for growth yield measurements.

**Figure S8:**
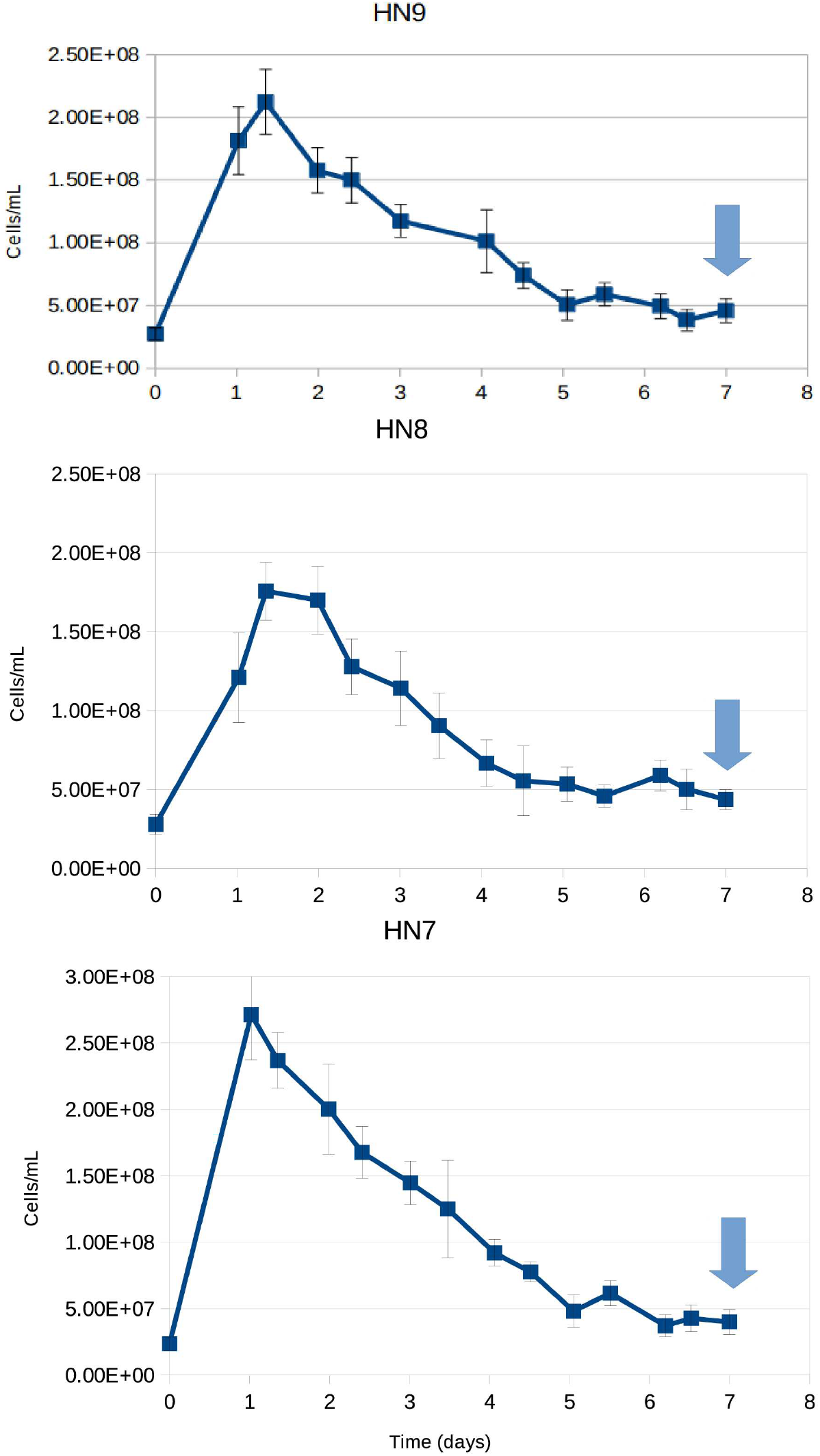
Growth curves for three replicate chemostat experiments grown with hydrogen and nitrate. Arrows indicate time of sampling for growth yield measurements.

**Figure S9:**
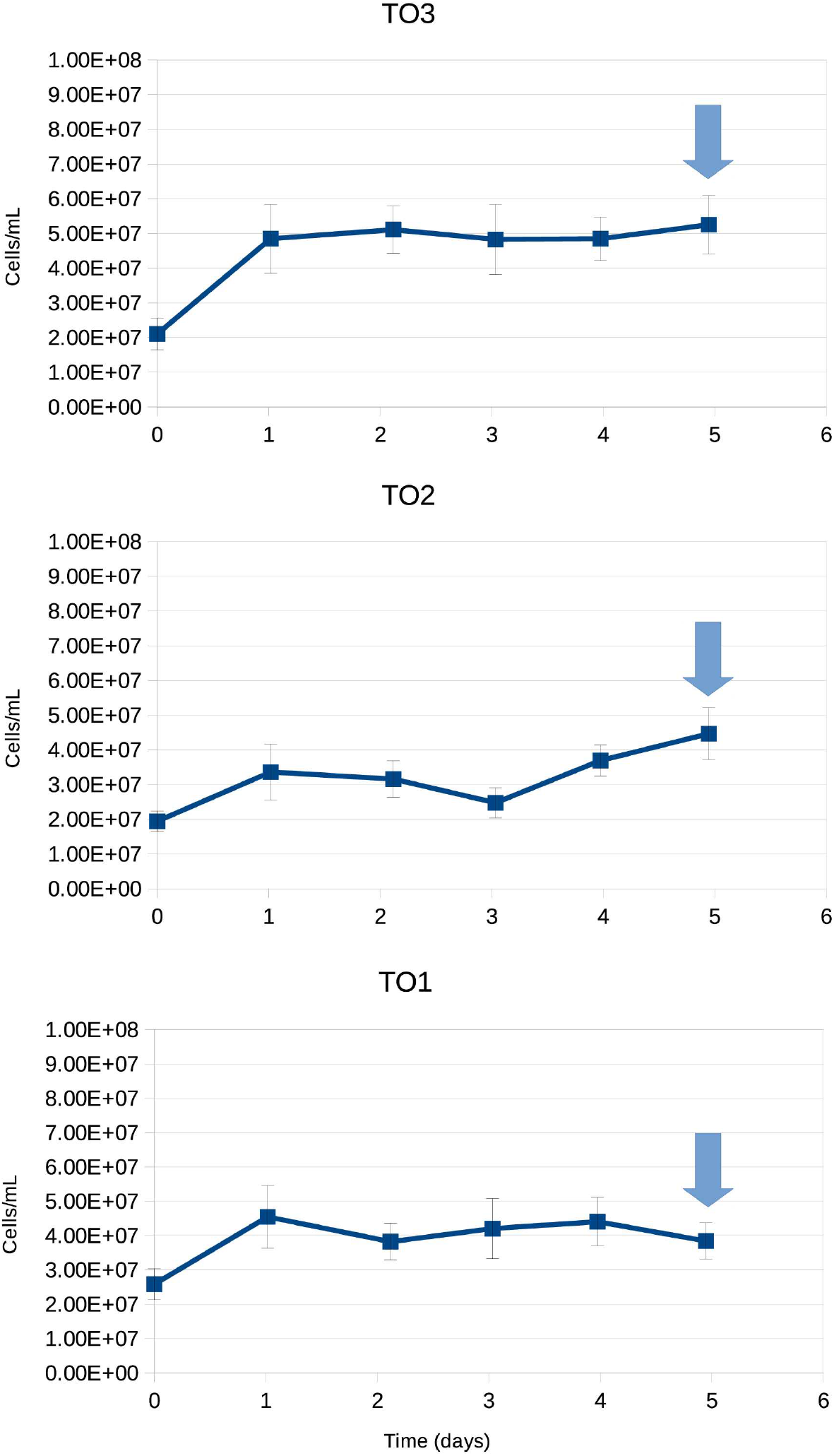
Growth curves for three replicate chemostat experiments grown with thiosulfate and oxygen. Arrows indicate time of sampling for growth yield measurements.

